# Regulation of *dctA* and DctA by cAMP-CRP and EIIA^Glc^ at the transcriptional and post-translational levels in *E. coli*: Consequences for aerobic uptake and metabolism of C_4_-dicarboxylates

**DOI:** 10.1101/2021.12.01.470772

**Authors:** Christopher Schubert, Gottfried Unden

**Author notes:** Address for correspondence: Christopher Schubert.

## Abstract

The expression of *dctA*, encoding the aerobic C_4_-dicarboxylate (C_4_-DC) transporter DctA of *Escherichia coli*, and its use in the presence of alternative carbon sources was characterized. *dctA* is regulated by cAMP-CRP and substrates that control cAMP levels, either through the phosphotransferase system (PTS), or through their metabolic link to PEP synthesis. The data indicates that phosphorylation of the regulator EIIA^Glc^ of the glucose-specific PTS represents the mediator for regulation. The *dctA* promotor region contains a class I CRP-binding site (position -81.5) and a DcuR-binding site (position -105.5). The response regulator DcuR of the C_4_-DC-activated DcuS-DcuR two-component system is known to stimulate expression of *dctA*, and cAMP-CRP is known to stimulate expression of *dcuS-dcuR*. Thus, activation of *dctA* expression by cAMP-CRP and DcuR is organized in a coherent feed-forward loop (FFL) where cAMP-CRP positively regulates the expression of *dctA* by direct stimulation and by stimulating the expression of *dcuR*. Stimulation by DcuR is presumed to require DNA bending by cAMP-CRP. In this way, CRP-FFL integrates carbon catabolite control and C_4_-DC-specific regulation. Moreover, EIIA^Glc^ of the glucose-specific PTS strongly interacts with DctA, which could lead to substrate exclusion of C_4_-DCs when preferred carbon substrates such as sugars are present. Since C_4_-DCs are perceived in the periplasmic space by the sensor DcuS, the substrate exclusion is not linked to inducer exclusion, contrasting classical inducer exclusion known for the lactose permease LacY. Thus, aerobic C_4_-DC metabolism is tightly regulated at the transcriptional and post-translational levels, whereas uptake of L-aspartate by DcuA is essentially unaffected. Overall, transcriptional and post-translational regulation of *dctA* expression and DctA function efficiently fine-tunes C_4_-DC catabolism in response to other preferred carbon sources.

## Introduction

DctA is a proton potential driven C_4_-dicarboxylate (C_4_-DC) transporter, which is responsible for the uptake of fumarate, succinate, L-, and D-malate, while serving as a coregulator of DcuS, under aerobic conditions (Kay and Kornberg, 1969; Kay and Kornberg, 1971; Janausch *et al*., 2002; Steinmetz *et al*., 2014). The transcription of *dctA* is controlled by the C_4_-DC two-component system DcuS-DcuR (Zientz *et al*., 1998; Davies *et al*., 1999). Furthermore, *dctA* is subject to transcriptional regulation by catabolite control and aerobic conditions by the cyclic AMP receptor protein (CRP) and the ArcBA two-component system (Davies *et al*., 1999; Gosset *et al*., 2004; Goh *et al*., 2005).

The phosphoenolpyruvate:glucose-specific phosphotransferase system (PTS) catalyzes the transport and phosphorylation of a variety of sugars and sugar derivatives. The PTS also mediates the regulation of carbon metabolism in glucophilic *E. coli*, where glucose is the preferred carbon source. The PTS transfers a phosphoryl group from the donor PEP via the EI, HPr and EIIA^Glc^ proteins to the glucose transporter EIIBC (Fig. 1). The phosphorylation status of EIIA^Glc^ depends on the availability of glucose and PEP. In the presence of glucose, phosphorylated EIIA^Glc^ transfers the phosphoryl group to EIIB, which phosphorylates glucose to glucose-6-phosphate during uptake by group translocation (Fig. 1). In the presence of glucose, EIIA^Glc^ is predominantly dephosphorylated and can inhibit transporters of alternative substrates that are subject to inducer exclusion such as the lactose permease LacY. In the absence of glucose, EIIA^Glc^ is primarily phosphorylated and stimulates cAMP production by the stimulation of adenylate cyclase (CyaA). The cAMP-CRP complex induces glucose repressed genes, such as the *lacZYA* operon for lactose catabolism. Sugars and sugar derivatives can be classified as PTS substrates, e.g., glucose, mannitol, and mannose, that are transported by PTS linked transporters, whereas non-PTS substrates are transported by primary or secondary transporters. In glycolytic bacteria such as *E. coli*, where the PTS system is the basis for glucose repression and carbon regulation, the PEP/pyruvate ratio is an important indicator of carbon metabolism (Deutscher *et al*., 2006). In addition, also non-PTS substrates, such as glycerol, maltose, and melibiose, that produce PEP during degradation, affect the phosphorylation state of EIIA^Glc^ and thus cAMP-CRP signaling (Hogema *et al*., 1998; Eppler and Boos, 1999; Eppler *et al*., 2002).

**Fig. 1:**
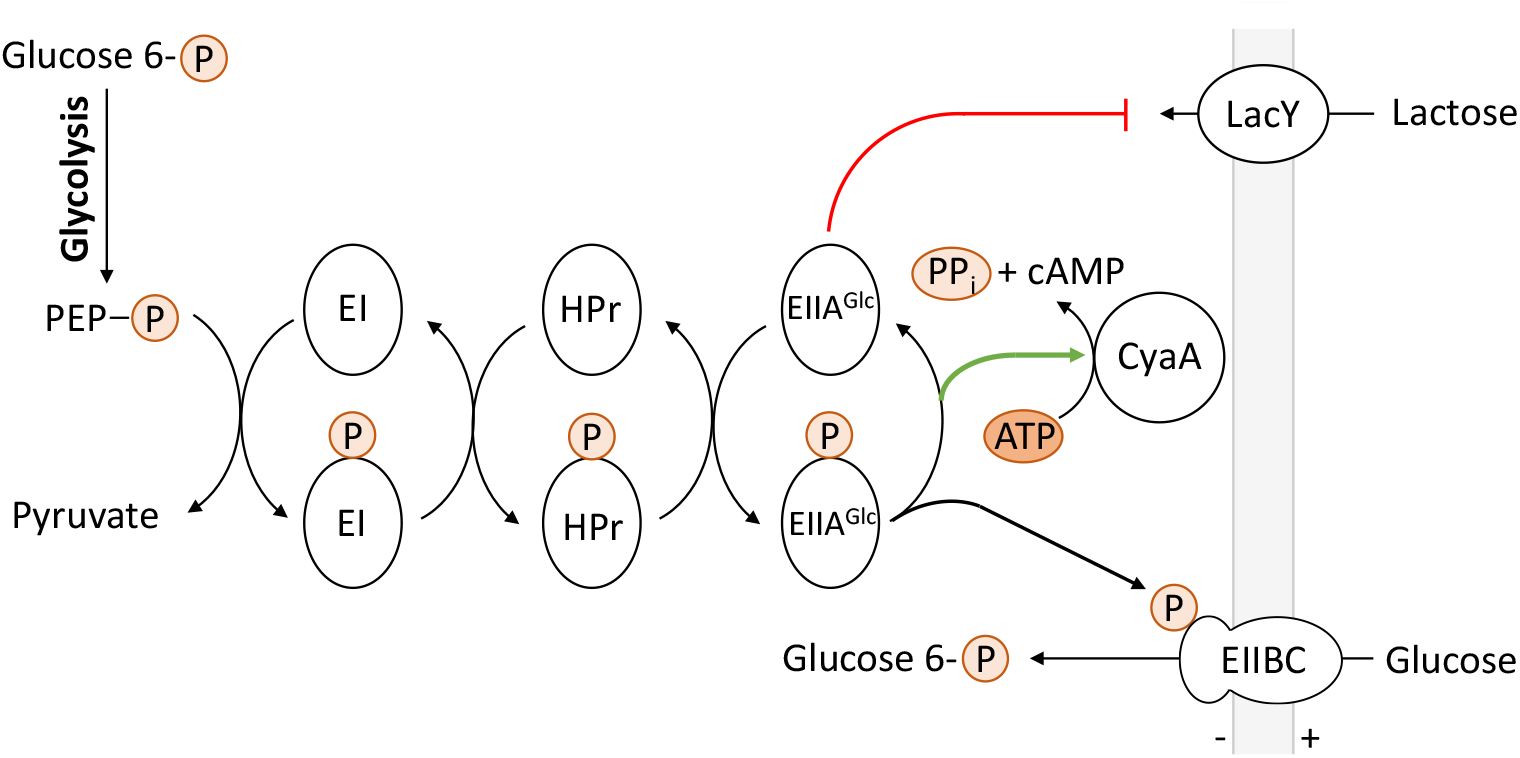
The phosphoenolpyruvate:glucose-specific phosphotransferase system PTS of *E. coli*. Abbreviations: PEP: phosphoenolpyruvate; EI: enzyme I *ptsI*; HPr: phosphocarrier protein HPr *ptsH*; EIIA^Glc^: enzyme IIA crr; EIIBC: glucose-specific PTS permease *ptsG*; P: phosphoryl group; CyaA: adenylate cyclase; LacY: lactose permease.

In general, catabolite regulation in *E. coli* is regulated by the EIIA/EIIA-P ratio, which is influenced by two factors: (I) Transport of the substrate by the PTS, and (II) influence of the substrate on PEP synthesis. (I) PTS substrates, in particular glucose, dephosphorylate EIIA^Glc^ during transport and thus inhibit the stimulation of cAMP production by the P-EIIA^Glc^-CyaA interaction. (II) The PEP/pyruvate ratio affects the PTS because PEP is the phosphoryl group donor for the EIIA^Glc^ phosphorylation cascade. Substrates such as C_4_-DCs that are channeled into central metabolism via the citric acid cycle (TCA) and without direct PEP production require (endergonic) reactions for PEP synthesis such as the reactions of gluconeogenesis. As a consequence, the PEP levels, and the rates of PEP synthesis on these substrates are low and tightly regulated (Gottschalk, 1986; Fuchs, 2014), resulting ultimately in low PEP availability (Hogema *et al*., 1998). C_4_-DCs, acetate and many amino acids do not produce PEP directly during degradation, require the reactions of gluconeogenesis and cause low PEP/pyruvate ratios. In the same way, substrates subject to inducer exclusion like lactose are not able to produce PEP under the respective conditions. In contrast, PTS substrates, which are usually sugars or sugar alcohols, directly produce PEP during degradation. The same is applicable to some other non-PTS substrates such as glycerol, which joins the glycolytic reactions upstream of PEP synthesis. In this work, a distinction is made between PTS and non-PTS substrates and substrates that are degraded with or without direct PEP synthesis during degradation.

CRP regulates the expression of more than 180 genes in *E. coli* (Grainger *et al*., 2005). Most of these genes are involved in catabolism of secondary (or non-PTS) carbon sources, such as lactose, glycerol, and maltose. Additionally, CRP is involved in a multitude of other processes, e.g., nitrogen assimilation (Mao *et al*., 2007), osmoregulation and virulence (Balsalobre *et al*., 2006). A recent protemic analysis identified metabolic and transcriptional regulation that is likely influenced by a network between C_4_-DC-specific (DcuS-DcuR) and carbon catabolite regulation (cAMP-CRP) (Surmann *et al*., 2020). CRP is activated by the secondary messenger cAMP and is subsequently able to bind to DNA (Deutscher *et al*., 2014). The homo-dimeric CRP belongs to the CRP-FNR superfamily of transcription factors (Green *et al*., 2001; Körner *et al*., 2003). The C-terminal domain of CRP carries a helix-turn-helix (HTH) motif for DNA-binding. Upon binding of cAMP, the C-terminal helix and the DNA-binding domain are elongated and reorganized, which results in uncovering the DNA recognition helix (Popovych *et al*., 2009; Sharma *et al*., 2009), followed by insertion of the HTH recognition helices into two adjacent DNA major grooves (Kim *et al*., 1992). cAMP-CRP causes 87° DNA bending by binding to the CRP consensus DNA binding site (Ebright *et al*., 1989; Parkinson *et al*., 1996). The nucleotide sequence and the location of the CRP-binding site are the key factors for determining gene expression. Promoter activation by cAMP-CRP can be clustered into three classes (Busby and Ebright, 1999; Fic *et al*., 2009). Class I promoters have a single CRP site located upstream of the RNA polymerase (RNAP) binding site and require only CRP to activate transcription. Class II promoters differ from class I promoters in that the CRP-binding site and the RNAP-binding site are shared (−35 promoter sequence). Class III promoters require additional transcriptional activators. The activating complex consists then of two or more cAMP-CRP complexes, or of one cAMP-CRP complex and an additional transcriptional activator that synergistically activate transcription (Fic *et al*., 2009).

In this work, the interaction between EIIA^Glc^ and DctA was studied by a bacterial two-hybrid system. In addition, the expression of *dctA* in response to different carbon sources in the context of cAMP-CRP activation was investigated. This work should expand the knowledge about the regulation of aerobic C_4_-DC metabolism at the transcriptional and post-translational levels, with DctA representing the main C_4_-DC transporter and thus the starting point for aerobic C_4_-DC degradation by *E. coli*.

## Results

### *dctA* expression is highly sensitive to numerous carbon sources

To study the response of the *dctA* promoter (*dctAp*) to different carbon sources, *dctAp* was genetically fused to the *lacZ* reporter gene, which encodes β-galactosidase. The *dctAp-lacZ* expression was examined in the presence of various pentoses, hexoses, disaccharides, sugar alcohols, and C_4_-DCs. Expression analysis of *dctAp-lacZ* was also performed in a *dctA*-deficient reporter strain to render DcuS constitutively active and thereby render *dctAp-lacZ* independent of transcriptional regulation by DcuS-DcuR (Davies *et al*., 1999; Janausch *et al*., 2002; Janausch *et al*., 2004; Kleefeld *et al*., 2009; Steinmetz *et al*., 2014). Bacteria were grown aerobically in LB medium in presence of different carbon sources. As expected (Davies *et al*., 1999; Steinmetz *et al*., 2014), *dctAp-lacZ* activity was higher (about five-fold) in the *dctA*-deficient reporter strain than in the wild type (Fig. 2A and B) and only the reporter strain with a wild-type *dctA* background shows a stimulation of *dctAp-lacZ* expression by C_4_-DCs compared to the control in LB medium (Fig. 2A). The substrates are ordered by their repression strength in Fig. 2. Starting with lactose, the expression of *dctAp-lacZ* decreased continuously, with glucose showing the highest repression compared to the control (Fig. 2A).

**Fig. 2:**
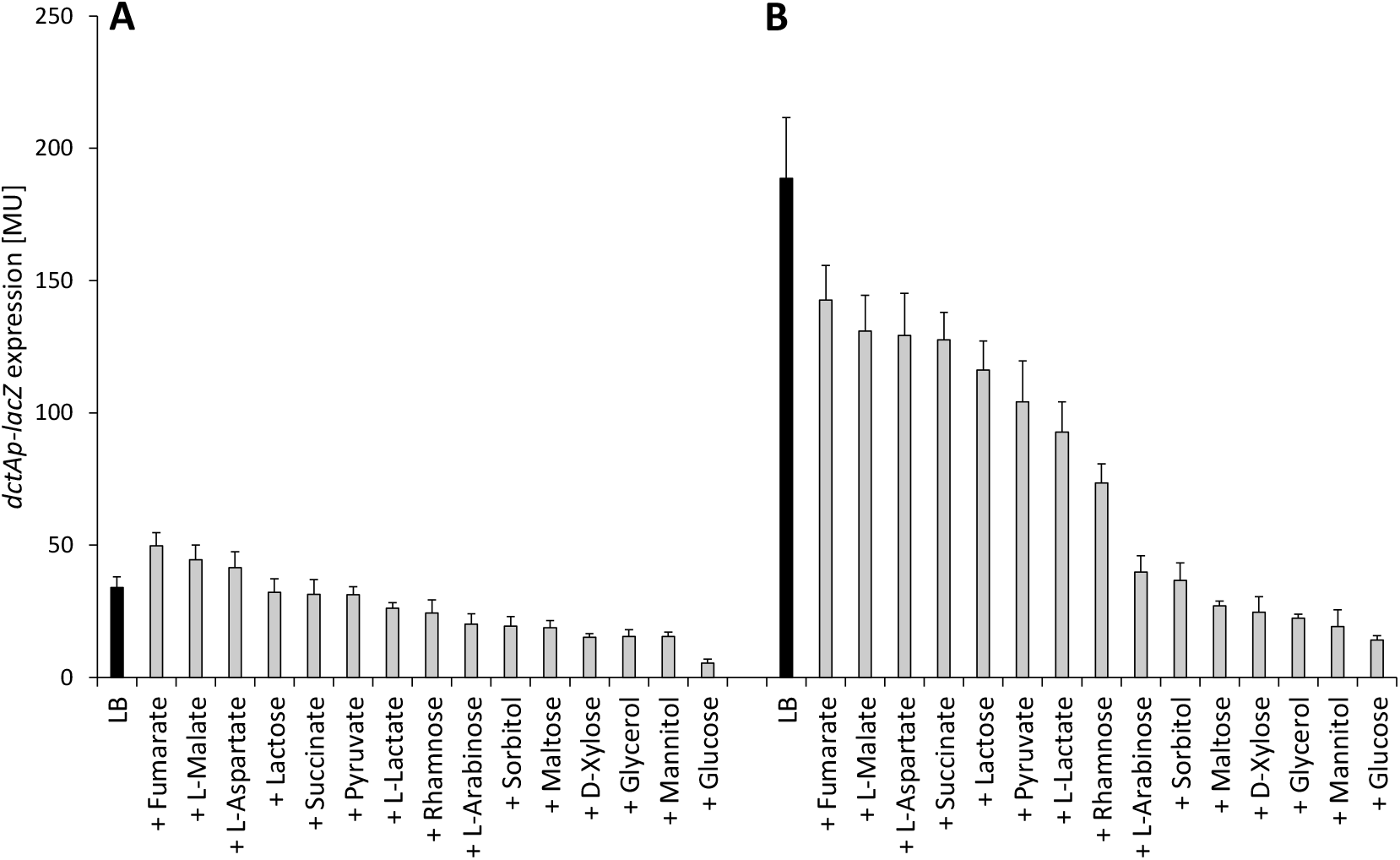
Expression of *dctAp-lacZ* in the presence of different carbon sources. The reporter strains (**A**) IMW385 (*dctAp-lacZ*, wild type *dctA*) and (**B**) IMW386 (*dctAp-lacZ*, Δ*dctA*) were cultivated aerobically at 37°C in LB medium in presence of different carbon sources (20 mM, gray bars). Control activities in absence of an effector are given (black bars). The β-galactosidase assay was performed in the late exponential growth phase (OD_578_= 1.5) and the β-galactosidase activity was quantified in Miller-Units (MU).

The *dctA*-deficient reporter strain that exhibits higher expression of the reporter, generally showed a higher repression of *dctAp-lacZ* by the substrates compared to the control (Fig. 2B). Moreover, even C_4_-DCs repress *dctAp-lacZ* compared to the LB reference (Fig. 2B). Notably, the order in which carbon sources repress *dctAp-lacZ* expression is quite similar in both strains (Fig. 2). The hexoses, glucose, and mannitol that are transported by the PTS showed the highest repression of *dctAp-lacZ*. Maltose, glycerol, and D-xylose stood out as non-PTS sugars by their high repression (Fig. 2). Both types of substrates (PTS and non-PTS) have in common that they directly produce PEP during degradation, in the case of L-arabinose and D-xylose via the pentose phosphate pathway (Mayer and Boos, 2005). These substrates are followed in their repression by pyruvate and rhamnose, and eventually C_4_-DCs, and the disaccharide lactose. The C_4_-di-/tricarboxylates and lactose showed the weakest repression of *dctAp-lacZ* (Fig. 2). Therefore, the response of *dctAp-lacZ* expression to various carbon sources is complex and obviously responds to factors related to CRP regulation such as the use of PTS/non-PTS pathways and direct production of PEP, in addition to specific regulation and inducer exclusion of the alternative substrates. Thus, the substrates with very high repression (glucose, sorbitol, and mannitol) are PTS substrates and directly produce PEP at a high rate. The next group with high repression is represented by non-PTS substrates (L-arabinose, maltose, D-xylose, and glycerol), which also produce PEP at a high rate. The composition of the low-repression substrate group is heterogeneous but is apparently characterized by the fact that no direct PEP production occurs during their degradation and the necessity of gluconeogenic reactions (C_4_-DCs, L-lactate, and pyruvate), or by additional regulation like inductor exclusion for lactose (Deutscher *et al*., 2006).

The effect of rhamnose is intermediate. Rhamnose is degraded to the C_3_-compounds dihydroxy acetone phosphate and lactaldehyde, and only degradation of the former directly delivers PEP.

Overall, there appears to be a direct relationship between *dctAp-lacZ* expression and metabolism of alternative substrates: PTS substrates and substrates that yield PEP at high rates during their degradation repress *dctA*, strongly suggesting a role for cAMP-CRP in regulating *dctA* expression. Furthermore, *dctAp-lacZ* highlights the ability of *E. coli* to continuously fine-tune gene expression in response to environmental factors, such as carbon sources.

### Effect of glycerol and pentoses on *dctAp-lacZ* expression

Analysis of *dctAp-lacZ* expression suggests an important role for PEP synthesis on the transcriptional regulation of the reporter fusion. The pentose D-xylose and the sugar alcohol glycerol showed strong repression of *dctAp* which can be explained by the efficient production of PEP from these substrates and correspondingly high PEP/pyruvate ratios (Fig. 3A). The effect of D-xylose and glycerol on *dctA* expression is further characterized in the following section to determine whether this observation is due to specific transcriptional regulation (GlpR or XylR) or carbon catabolite regulation by cAMP-CRP (Fig. 3A). Glycerol uptake is catalyzed by GlpF and phosphorylation by GlpK, producing glycerol-3-phosphate (G3P). Glycerol dehydrogenase GlpD oxidizes G3P to dihydroxyacetone phosphate (DHAP), and DHAP is fed into glycolysis with direct synthesis of PEP (Schryvers *et al*., 1978; Yeh *et al*., 2008). D-Xylose enters the cell either through proton motive force (XylE) or an ATP-driven (XylFGH) transporter. The isomerase XylA converts D-xylose to D-xylulose and subsequently the xylulokinase XylB produces D-xylulose 5-phosphate (X5P), an intermediate of the pentose phosphate pathway (Mayer and Boos, 2005). The oxidative pentose phosphate pathway produces hexose-P and glyceraldehyde-3-P which are degraded by glycolysis with efficient production of PEP.

**Fig. 3:**
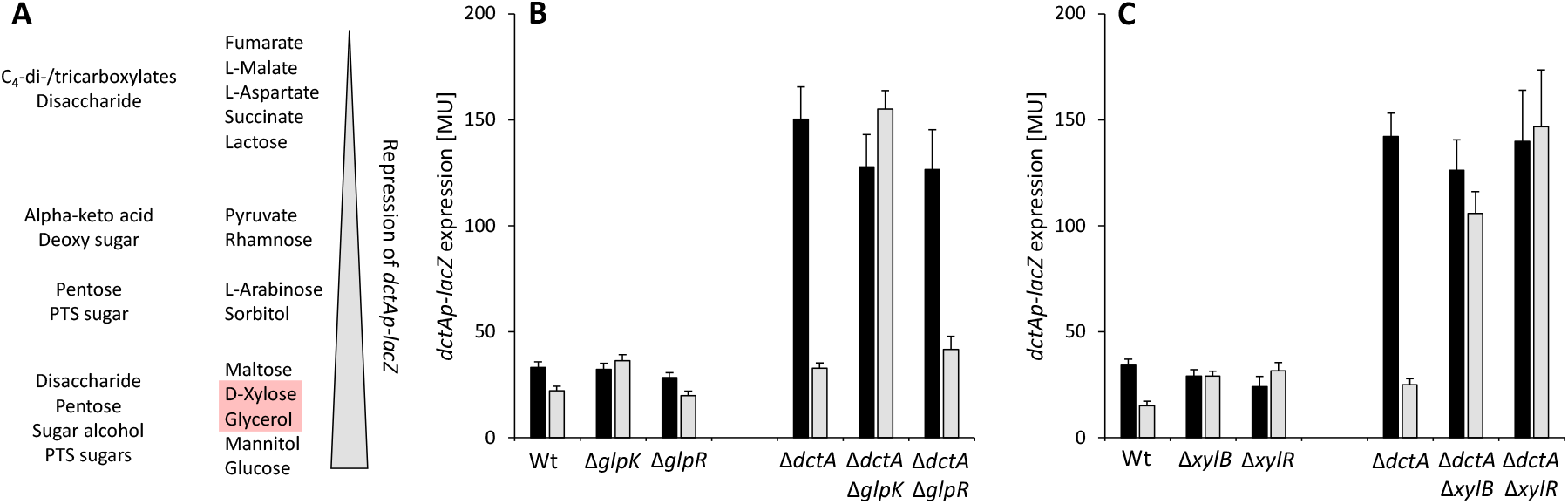
*dctAp-lacZ* expression in mutants deficient in glycerol (B) and D-xylose (C) metabolism. (**A**) All tested carbon sources were sorted according to the repression strength of *dctAp-lacZ* expression. The substrate class is described on the left side. Bacteria containing the *dctAp-lacZ* fusion were cultivated aerobically at 37°C in LB medium (black bars) and in presence of glycerol (**B**) or D-xylose (**C**) (20 mM, gray bars) to the late exponential growth phase (OD_578_= 1.5). The β-galactosidase activity was quantified in Miller-Units (MU). Abbreviations: GlpK: glycerol kinase; GlpR: transcriptional repressor; XylB: xylulokinase; XylR: transcriptional dual regulator.

To test whether these pathways are involved, the corresponding kinases (*glpK* and *xylB*) and transcriptional regulators (*glpR* and *xylR*) were genetically inactivated in both *dctAp-lacZ* reporter strains. Bacteria were cultivated aerobically in LB medium with either glycerol or D-xylose as an effector.

The *dctA* repression by glycerol was abolished in the *glpK*-deficient strain (Fig. 3B). GlpK catalyzes the phosphorylation of glycerol to G3P, whereas the specific regulator GlpR of glycerol metabolism had no effect on *dctAp-lacZ* expression (Fig. 3B). These observations are consistent with the assumption that high PEP levels derived from glycerol cause the effect on *dctA* expression, confirming similar observations made for *malTp-lacZ* expression (Eppler and Boos, 1999). Accordingly, the repression of *malT* by glycerol was shown to depend on cAMP-CRP (Eppler and Boos, 1999), presumably also via PEP and its effect on cAMP-CRP levels. A similar observation was made for D-xylose, where genetic inactivation of the xylulokinase XylB resolved *dctA* repression. Interestingly, the XylR-deficient mutant showed the same effect (Fig. 3C). This observation can be explained by a previous work that showed how *xylR*-deficiency abolished the expression of the *xylAB* and *xylFGHR* operons (Song and Park, 1997), suggesting that this phenotype is due to the inability of the *xylR*-deficient strain to express XylB. This would suggest X5P-specific repression, as previously seen with G3P, which feeds into glycolysis and produces PEP. Either way, the observed repression of *dctAp-lacZ* by glycerol and D-xylose is likely attributable to the efficiency of PEP conversion and the resulting effects on cAMP-CRP levels.

### *dctA* expression depends on cAMP-CRP activation

cAMP-CRP is essential for the activation of catabolite-controlled genes other than glucose (Kolb *et al*., 1993; Deutscher *et al*., 2006; Fic *et al*., 2009). The PEP/pyruvate ratio indirectly affects cAMP production via the PTS for which PEP is the phosphoryl group donor. Substrates leading directly to PEP synthesis in catabolism have a higher PEP/pyruvate ratio than substrates without direct PEP production which affects the phosphorylation state of EIIA^Glc^ and thus cAMP production by stimulation of the adenylate cyclase CyaA (Hogema *et al*., 1998). Accordingly, cAMP levels fluctuate depending on transport (PTS vs. non-PTS substrate), metabolic link to PEP production (Bennett *et al*., 2009), and additional individual regulation. Therefore, cAMP-CRP is able to fine-tune regulation of genes of alternative metabolic pathways in response to available carbon sources.

The adenylate cyclase CyaA was genetically inactivated in the *dctAp-lacZ* reporter strains, in order to impair activation by cAMP-CRP (Fig. 4). Expression of *dctAp-lacZ* decreased 6.3- and 6.2-fold in the presence of glucose or the *cyaA* mutant, respectively (Fig. 4A), whereas under *dctA*-deficiency, *dctAp-lacZ* activity decreased 13.4- and 13.8-fold (Fig. 4B). Therefore, the absence of DctA results in a significantly stronger reduction in the CyaA-deficient mutant, confirming that DcuR and cAMP-CRP regulate the expression of *dctA*. However, C_4_-DC-dependent induction by DcuS-DcuR is inferior to activation by cAMP-CRP and occurs only when sufficient amounts of cAMP-CRP complexes are present. Overall, transcription of *dctA* is highly dependent on cAMP-CRP activation in response to various carbon sources, whereas C_4_-DC-specific stimulation by DcuS-DcuR is inferior and requires the presence of cAMP-CRP.

**Fig. 4:**
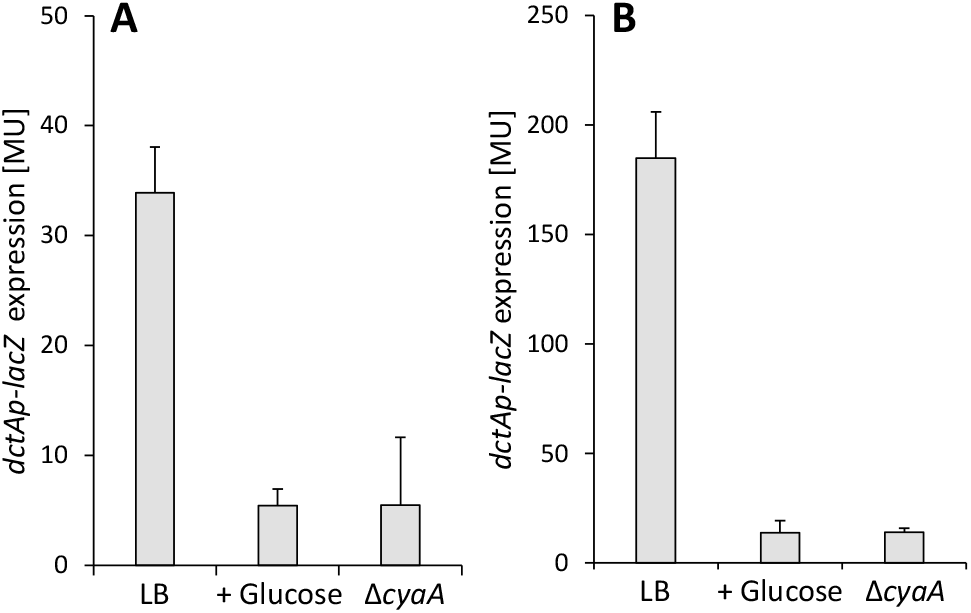
Dependence of *dctAp-lacZ* on cAMP-CRP activation. Bacteria were cultivated aerobically at 37°C in LB medium and glucose (20 mM) to the late exponential growth phase (OD_578_= 1.2). β-Galactosidase activity was quantified in Miller-Units (MU). (**A**) IMW385 (*dctAp-lacZ*) and IMW669 (IMW385, Δ*cyaA*); (**B**) IMW386 (*dctAp-lacZ*, Δ*dctA*) and IMW665 (IMW386, Δ*cyaA*).

### Interaction between DctA and EIIA^Glc^ of the PTS

Bacteria producing the aerobic C_4_-DC transporter DctA and the protein EIIA^Glc^ from the glucose-specific PTS were tested for interaction using the bacterial two-hybrid system BACTH. The N-and C-termini of DctA have cytoplasmic location (Witan *et al*., 2012). Fusions of DctA to the T18 and T25 fragments of *Bordetella pertussis* adenylate cyclase were produced in all combinations. In the same way, the N- and C-termini of EIIA^Glc^ were fused genetically to the T18 and T25 fragments. When DctA and EIIA^Glc^ fused to T18 and T25, respectively, were produced in the reporter strain, high β-galactosidase activity was observed at the same level as in the positive control (_T18_Zip × _T25_Zip) or higher, clearly exceeding the background activity (_T18_Zip × DctA_T25_) (Fig. 5). The data indicates distinct interaction between DctA and EIIA^Glc^.

**Fig. 5:**
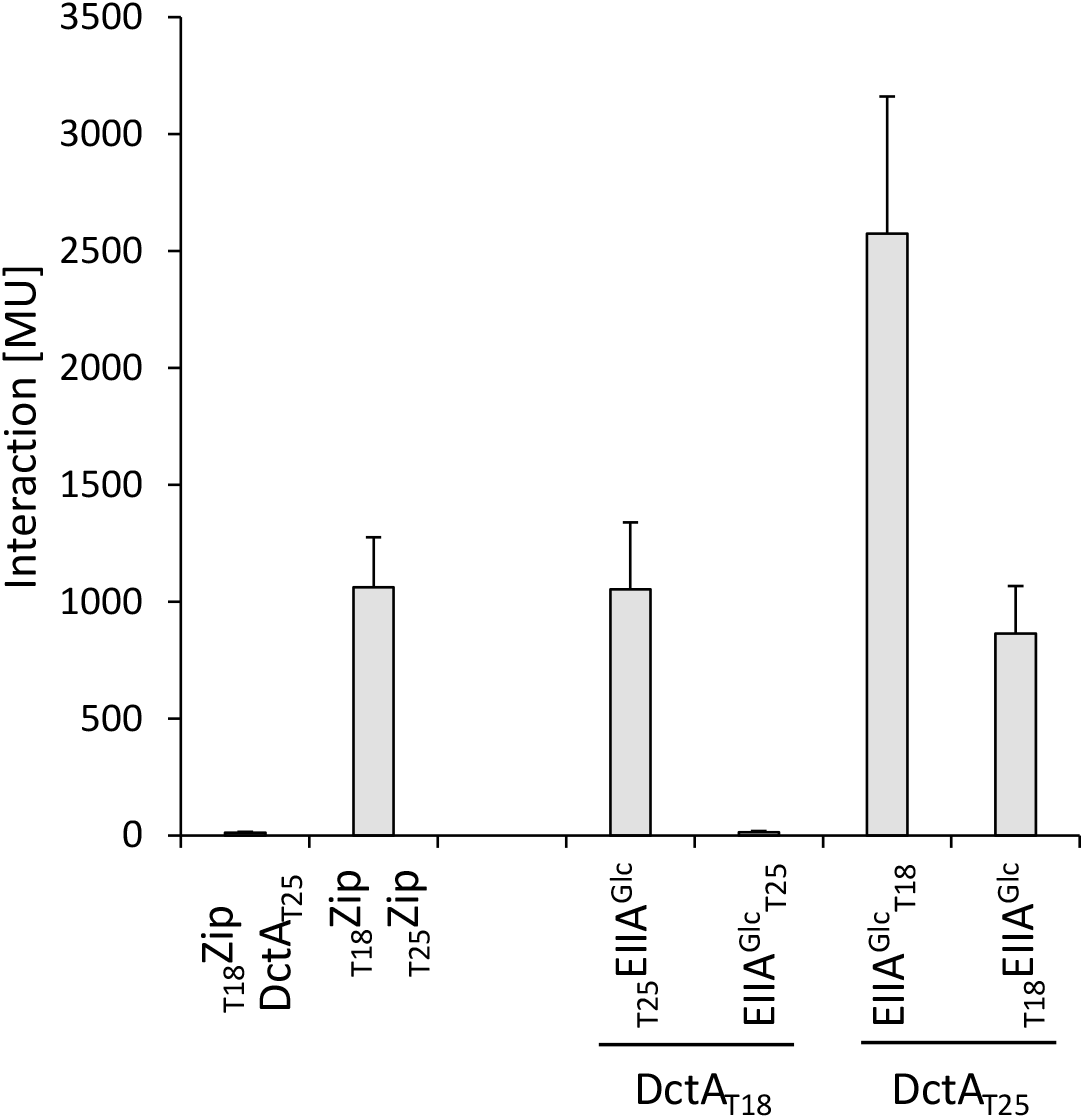
DctA-EIIA^Glc^ interaction *in vivo* using the BACTH system. *E. coli* BTH101(Δ*cyaA*) was co-transformed pairwise with plasmids encoding fusions of T25 to DctA/EIIA^Glc^ (_T25_DctA) and fusions of T18 to DctA/EIIA^Glc^ (_T18_DctA). The combinations are shown on the x-axis. The leucine zipper pair _T18_Zip and _T25_Zip are applied as positive control (Karimova *et al*., 1998; Karimova *et al*., 2001), the pair _T18_Zip/_T25_DctA as the negative control for background β-galactosidase activity. The corresponding plasmids are derivatives of pUT18C (_T18_EIIA^Glc^), pUT18 (DctA_T18_), pKT25 (_T25_EIIA^Glc^), and pKNT25 (DctA_T25_) (Tab. 1). The strains were grown aerobically in LB medium at 30°C to the exponential growth phase (OD_578_ = 0.5). β-Galactosidase activity was quantified in Miller-Units (MU).

## Discussion

*E. coli* can grow on a mixture of sugars and other carbon sources but consumes the available substrates in sequential order through a process known as catabolite repression (CR) (Görke and Stülke, 2008). The order depends, among other things, on the energy yield or the metabolic suitability of the substrate for *E. coli* (Stülke and Hillen, 1999; Leuze *et al*., 2012). This ensures that bacterial cells adapt their metabolic resources to the preferred carbon source, which in turn optimizes the growth rate. CR is a central regulatory mechanism that is involved in regulation of 5-10% of all genes (Stülke and Hillen, 1999; Brückner and Titgemeyer, 2002; Görke and Stülke, 2008; Aidelberg *et al*., 2014). The classic example of CR is the biphasic growth (diauxie) of *E. coli* on a glucose and lactose-containing medium, with the bacteria initially using glucose and then lactose (Monod, 1941; Loomis and Magasanik, 1967). In the absence of PTS substrates and availability of substrates causing a high PEP/pyruvate ratio, EIIA^Glc^ is phosphorylated and stimulates cAMP production by CyaA. The increase in intracellular cAMP leads to the formation of cAMP-CRP complexes and subsequent induction of genes requiring transcriptional activation by cAMP-CRP. Non-PTS substrates, such as glycerol, maltose, and melibiose, produce also high levels of PEP, affecting the phosphorylation state of EIIA^Glc^ and cAMP production (Hogema *et al*., 1998; Eppler and Boos, 1999; Eppler *et al*., 2002). These regulatory features highlight the ability of cAMP-CRP to indirectly sense carbon sources and thus modulate transcriptional regulation.

### The promoter of *dctA* represents a class I CRP-dependent promoter

Class I CRP-dependent promoters require cAMP-CRP for transcriptional activation. This type was identified for the *dctAp* reporter, which depends on cAMP-CRP activation and contains a CRP-binding site at a position characteristic of class I promoters. At Class I promoters the CRP dimer interacts with the α-C-terminal domain of the RNA polymerase (RNAP) (Zhou *et al*., 1994), which increases the affinity of RNAP for promoter DNA (Igarashi and Ishihama, 1991; Ebright, 1993; Savery *et al*., 1996) by proper positioning for open complex formation (Browning *et al*., 2019). cAMP-CRP induces a nucleic acid bend of 87° at the CRP consensus sequence (Ebright *et al*., 1989; Parkinson *et al*., 1996), but the degree of the bend depends on the nucleotide sequence of the CRP-binding site. The CRP and DcuR-binding sites of *dctA* are located at -81.5 and -105.5 upstream of the transcription start, respectively (Fig. 6A) (Davies *et al*., 1999; Abo-Amer *et al*., 2004). Expression and positioning data suggest that DcuR induction depends on cAMP-CRP-induced DNA bending, which places DcuR in proximity to RNAP (Fig. 7B).

**Fig. 6:**
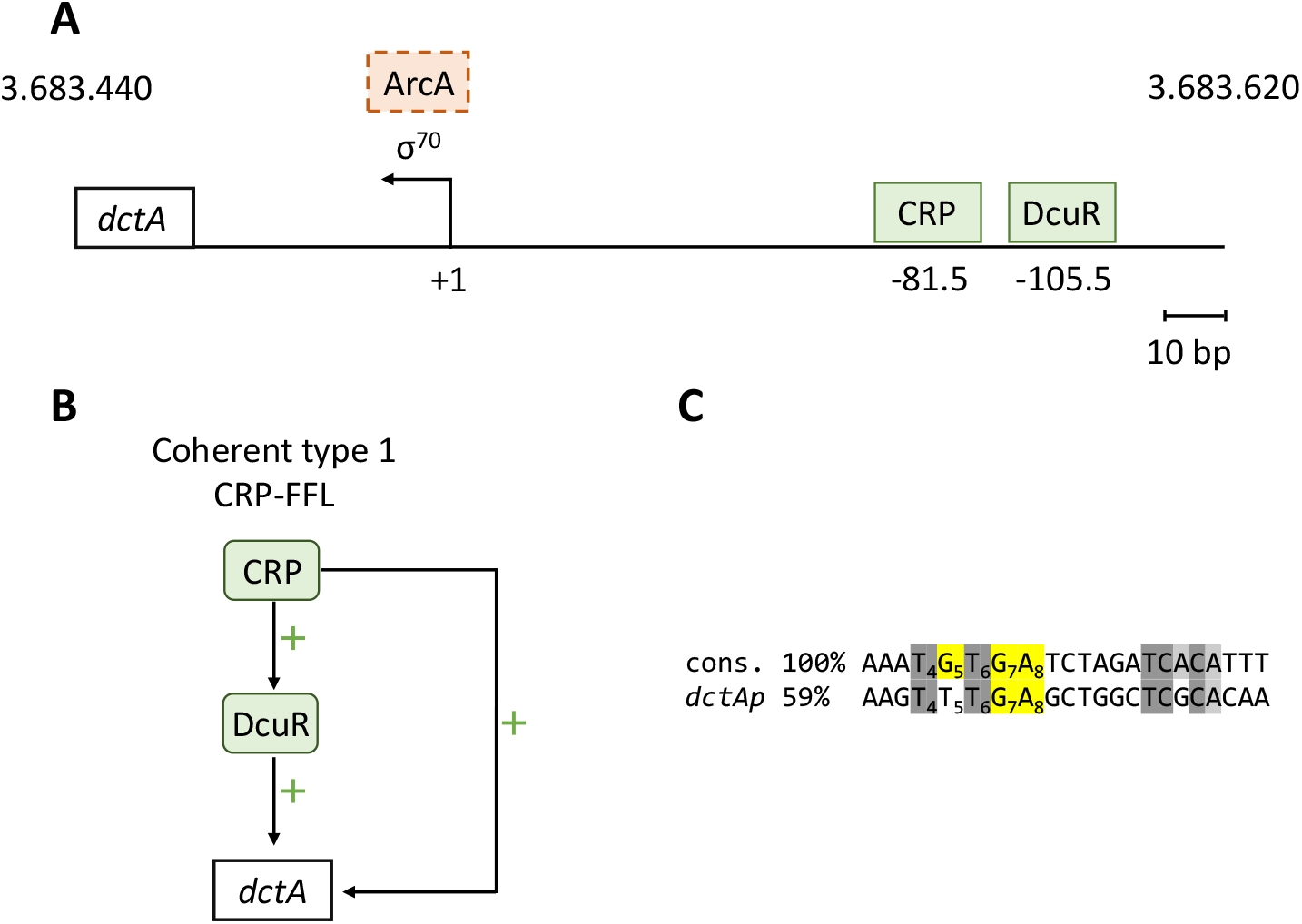
Annotation of the *dctA* promoter region. (**A**) Promoter region of *dctA*, including binding sites based on experimental data (CRP: Davies *et al*., 1999 and DcuR: Abo-Amer *et al*., 2004). (**B**) Feed-forward loops (FFL) of *dctA* regulation. (**C**) Alignment of the *dctAp* CRP-binding site with the CRP consensus (cons.) (Ebright *et al*., 1989). CRP-binding sites were acquired via RegulonDB (Gama-Castro *et al*., 2016). The nucleotides in direct contact with the CRP protein are annotated (yellow) (Leuze *et al*., 2012).

**Fig. 7:**
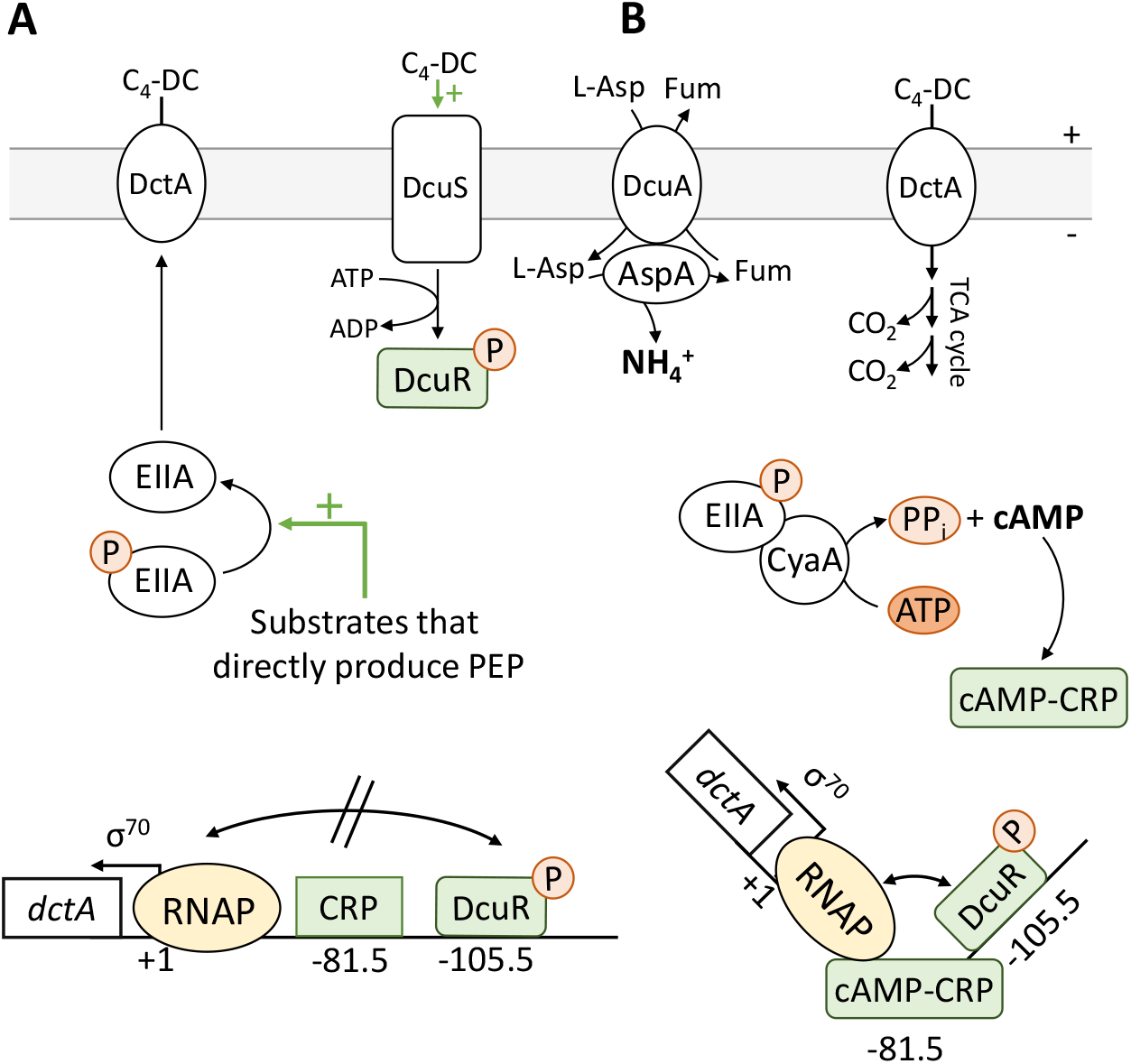
Model for the regulation of C_4_-dicarboxylate metabolism during aerobic growth. C_4_-dicarboxylate metabolism in aerobic growth (**A**) in presence and (**B**) in absence of a substrate upstream of PEP synthesis. Abbreviations: DctA: aerobic C_4_-dicarboxylate transporter; DcuA: aerobic L-Asp transporter; C_4_-DC: C_4_-dicarboxlate; RNAP: RNA-polymerase; CyaA: adenylate cyclase; EIIA: enzyme IIA^Glc^; PEP: phosphoenolpyruvate; DcuS-DcuR: C_4_-DC-specific two-component system.

Glycerol has an impact on transcriptional regulation via cAMP-CRP (Hogema *et al*., 1998). This observation was further illustrated by the *dctAp-lacZ* reporter gene assays (Fig. 2). Proteomic data from *E. coli* cultured under different carbon sources in minimal M9 medium showed CRP at a rather constant level of 2943 (±535 18,17%) copies per cell (Schmidt *et al*., 2016). The cAMP concentrations varied widely in the presence of glucose (35 μM), glycerol (83 μM 137% increase), and acetate (146 μM 317% increase) (Bennett *et al*., 2009), confirming that essentially the intracellular cAMP concentrations determine cAMP-CRP activation. In addition, the quality and quantity of the CRP-binding site is also critical. Quality refers to the nucleotide sequence of the CRP-binding site, whereas quantity reflects the frequency of CRP-binding sites before a transcription start.

### The CRP-binding site of *dctA* differs compared to the CRP consensus

The consensus sequence of CRP-binding sites (5′-AAA**TGTGA**TCTAGA**TCACA**TTT-3′) is palindromic, with the consensus half site 5′-A_1_A_2_A_3_**T**_4_**G**_5_**T**_6_**G**_7_**A**_8_T_9_C_10_T_11_-3’ (Busby and Ebright, 1999; Leuze *et al*., 2012) (Fig. 6C). The first half of the binding site is dominant in TF binding and therefore conserved. CRP directly interacts with G_5_, G_7_ and A_8_ in the core motif of the first half site (Schultz *et al*., 1991; Chen *et al*., 2001; Leuze *et al*., 2012). Mutations in the CRP site towards the consensus lead to an enhanced ability of cAMP-CRP to activate the promoter and thereby reprogram the hierarchy in sequential carbon utilization (Narang *et al*., 1997; Kovárová-Kovar and Egli, 1998; Aidelberg *et al*., 2014). Interestingly, G_5_ of the first half site of the CRP-binding site in *dctAp* is changed to T_5_, which likely weakens the interaction with cAMP-CRP (Fig. 6C). Variations in the second half of the CRP-binding site are less important for cAMP-CRP interaction (Schultz *et al*., 1991; Chen *et al*., 2001; Leuze *et al*., 2012).

Besides the quality of the CRP-binding site, the total number of CRP-binding sites are an important characteristic in gene regulation (Leuze *et al*., 2012). The mannitol-specific PTS enzyme II MtlA transports and phosphorylates mannitol. Mannitol-1-phosphate 5-dehydrogenase MtlD converts mannitol-1-phosphate to fructose-6-phosphate, which enters glycolysis (Mayer and Boos, 2005). The *mtlAD* operon contains five CRP-binding sites (Ramseier and Saier Jr, 1995) with a conservation level between 58% and 75% compared to the CRP consensus (Ebright *et al*., 1989). However, a large fraction of CRP-controlled genes has only one CRP-binding site similar to *dctA* (Gama-Castro *et al*., 2016). Overall, *dctA* is a unique example of cAMP-CRP activation, which emphasize regulatory fine-tuning in response to available carbon sources and C_4_-DCs, combining regulation by global and specific transcriptional regulators, including cAMP-CRP and DcuS-DcuR.

### *dctA* is controlled by a CRP-FFL coherent type 1

Feed-forward loops (FFL) are important regulatory networks in *E. coli* that contain two transcription factors (TF) and one or more genes. In FFLs, TF1 directly controls the expression of the target gene and TF2, while TF2 also regulates the target gene, resulting in indirect regulation of the target gene by TF2 via TF1. In FFLs, a distinction is made between coherent and incoherent types. If the direct effect (positive or negative) of TF1 on the gene is the same as the indirect effect of TF1 via TF2, the FFL is termed coherent (Shen-Orr *et al*., 2002; Mangan and Alon, 2003; Yang *et al*., 2018). In the *dctA* feed-forward-loop (Fig. 6B), cAMP-CRP directly activates *dctA* and *dcuR* transcription (Davies *et al*., 1999; Oyamada *et al*., 2007), and DcuR positively regulates *dctA* (Abo-Amer *et al*., 2004). Thus, this network motif is characterized as a CRP-FFL of coherent type 1 (Fig. 6B) (Yang *et al*., 2018). This CRP-FFL indicates that sufficient cAMP-CRP must be present for specific transcriptional regulation to occur. This adds a second layer to the regulation of *dctA*. For example, in the presence of C_4_-DCs and glucose, glucose causes low levels of cAMP, preventing transcriptional activation of *dcuR* and *dctA* by cAMP-CRP. With C_4_-DCs as the carbon source, the cAMP levels are high, and cAMP-CRP activates transcription of *dcuR* and *dctA*, allowing specific transcriptional activation by DcuS-DcuR in response to C_4_-DCs.

### DctA and EIIA^Glc^ of PTS interact: Inhibition of DctA and substrate (C_4_-DC) exclusion by EIIA^Glc^?

*dctA* is tightly regulated by cAMP-CRP in response to carbon availability. The PTS network has also the ability to post-translationally inhibit transporters of secondary metabolic pathways via EIIA^Glc^. DctA and EIIA^Glc^ showed an interaction in the BACTH system, suggesting an interaction similar to LacY-EIIA^Glc^, causing inhibition of LacY and thus inducer exclusion (Dills *et al*., 1982). Although interaction of DctA with EIIA^Glc^ is strong and evident, the role and physiological consequence is not clear. The interaction is suggested to inhibit DctA and C_4_-DC uptake in the presence of the preferred substrate glucose (i.e., by dephosphorylated EIIA^Glc^) to subsequently exclude C_4_-DCs as alternative substrates from the cell. Inhibition of C_4_-DC uptake does not result in inducer exclusion, since C_4_-DCs are sensed in the periplasm by the two-component system DcuS-DcuR (Pappalardo *et al*., 2003; Unden *et al*., 2016). The supposed model suggests ‘substrate exclusion’ which is at variance with classical inducer exclusion, exemplified by the inhibition of lactose permease LacY or the maltose ABC transporter (Dills *et al*., 1982; Misko *et al*., 1987; Dean *et al*., 1990). Classical inducer exclusion inhibits uptake of substrates that serve also as inducers in the cytoplasm, thereby inhibiting the induction of alternative metabolic pathways by specific transcriptional regulation (Deutscher *et al*., 2006). The interaction between DctA and EIIA^Glc^ indicates a further level of regulation on DctA, in addition to the transcriptional regulation of *dctA* by cAMP-CRP (Davies *et al*., 1999).

### DctA is regulated by the PTS at the transcriptional and posttranslational levels

Expression and interaction analysis suggest that DctA is regulated by the glucose-specific PTS at the transcriptional and posttranslational levels centered on EIIA^Glc^ and cAMP-CRP. Under aerobic conditions, DcuA catalyzes an L-Asp:C_4_-DC (primarily fumarate) antiport, while AspA converts L-Asp to fumarate and ammonium (Strecker *et al*., 2018; Schubert *et al*., 2020). Ammonium is assimilated via the GS-GOGAT pathway and provides a nitrogen source that stimulates the nitrogen demand of *E. coli* (Schubert *et al*., 2020). EIIA^Glc^ interacts with DctA, suggesting that in the presence of substrates that directly yield PEP, unphosphorylated EIIA^Glc^ inhibits C_4_-DC uptake by DctA (Fig. 7A). As a consequence, the cAMP levels are low, resulting in basal expression of *dctA*, whereas DcuR is phosphorylated but fails to stimulate expression of *dctA* due to the low levels of cAMP-CRP (Fig. 7A). Basal expression of *dctA* is essential for the formation of the DctA-DcuS sensor complex, as this interaction converts DcuS to the C_4_-DC-responsive state (Wörner *et al*., 2017). In the absence of substrates that directly feed into PEP synthesis, EIIA^Glc^ is predominantly phosphorylated, which stimulates cAMP production by CyaA (Fig. 7B). The inhibition of DctA would be relieved and C_4_-DCs enter the bacterial cell. Moreover, cAMP-CRP and phosphorylated DcuR stimulate the expression of *dctA* (Fig. 7B). C_4_-DCs are introduced into the TCA cycle and oxidized to CO_2_. Under both conditions, the nitrogen shuttle catalyzed by DcuA-AspA is active and supplies the cell with ammonium (Fig. 7) (Schubert *et al*., 2020; Schubert *et al*., 2021).

C_4_-DCs have a comparatively low energy yield and require gluconeogenetic reactions which slows down the growth rate. Therefore, DctA is integrated into the PTS network on a transcriptional and post-transcriptional level to control aerobic C_4_-DC metabolism and its relation to substrates producing directly PEP without the need for gluconeogenetic reactions. The need for gluconeogenesis leads to significant changes in the proteome of *E. coli* after switching to C_4_-DCs as the main carbon source (Surmann *et al*., 2020). DcuA, as the main L-Asp transporter, appears to be independent of post-translational regulation. The nitrogen shuttle of DcuA and AspA continuously supplies *E. coli* with ammonium to saturate the nitrogen demand during growth (Strecker *et al*., 2018; Schubert *et al*., 2020). Tight regulation of DctA limits the reflux of C_4_-DC secreted by DcuA-AspA. On the one hand, DctA is necessary for the DcuS-DctA sensor complex, but on the other hand, uptake of C_4_-DCs is undesirable when better substrates are available. These adverse needs for C_4_-DC transport can be solved by the integration of DctA into the PTS network to fine-tune aerobic C_4_-DC metabolism to the availability of other carbon sources.

## Materials and Methods

### Bacterial strains and growth conditions

The *E. coli* K12 strains are listed in Table 1. All molecular methods, including phage P1 transduction, were performed according to standard procedures (Jones and Gunsalus, 1987; Miller, 1992; Kleefeld *et al*., 2009; Monzel *et al*., 2013). Phage P1 transduction to obtain *dctAp-lacZ* mutants were performed as described previously (Sambrook *et al*., 2001; Miller, 1992). The recipient strain was either IMW385(*dctAp-lacZ*) or IMW386(*dctAp-lacZ, dctA*::Spc^R^) and the donor strains were from the Keio collection (Baba *et al*., 2006). The success of recombination was verified by sequencing the corresponding region or growth on selective agar plates. Bacteria were grown aerobically at 37 °C in lysogeny broth (LB) as indicated. All media were inoculated at 37 °C with 1–5% (v/v) of an overnight culture grown under the same conditions and in the same medium.

**Tab. 1:**
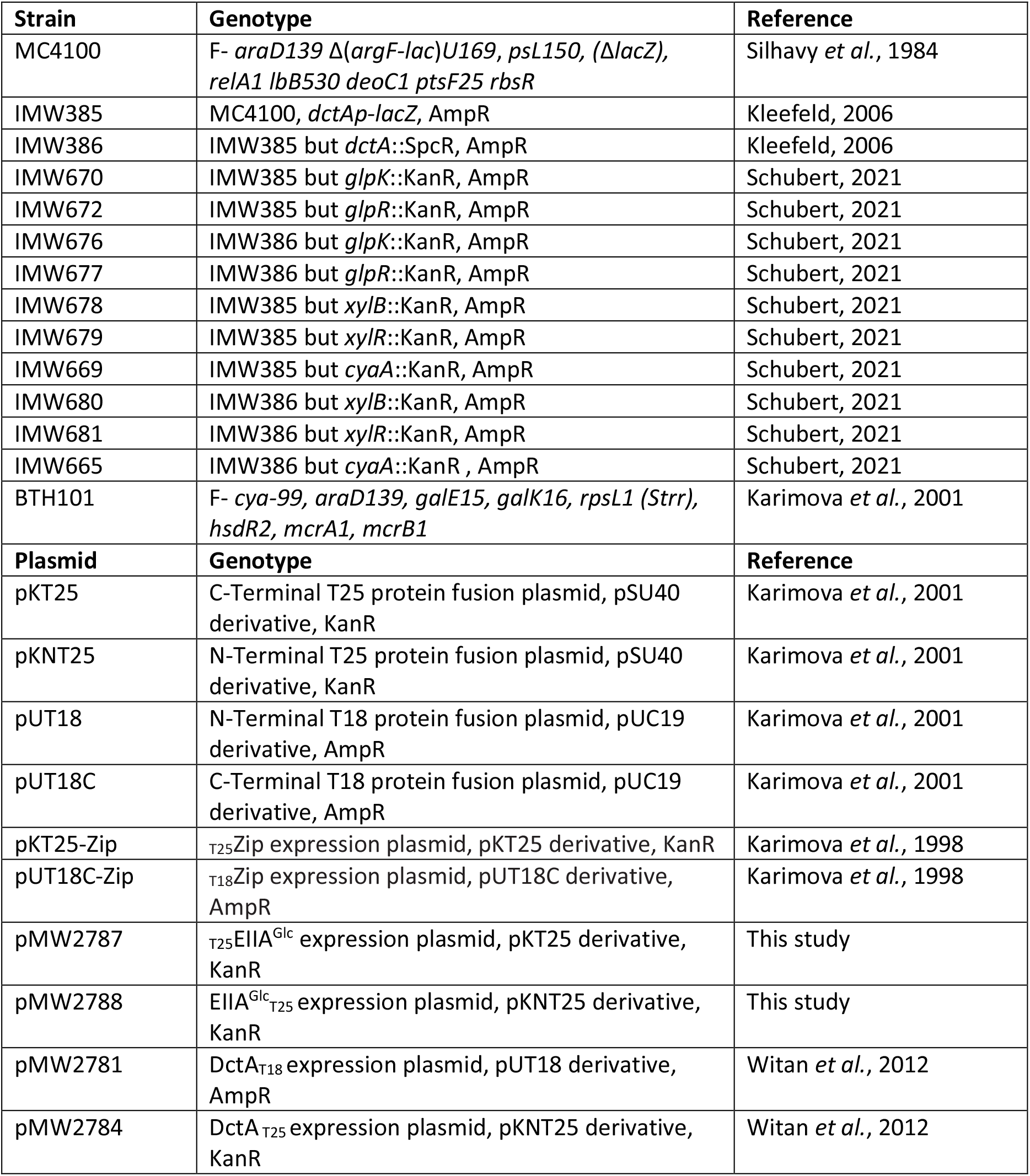
Strains of *E. coli* and plasmids used.

### β-galactosidase assay

Bacteria (Table 1) were grown aerobically in LB medium with effectors as stated in the experiment. Interactions in the BACTH system were measured in terms of β-galactosidase activity (Miller, 1992). BACTH experiments were conducted as described previously (Monzel *et al*., 2013) with slight modifications (Wörner *et al*., 2016). Activities are the mean of at least two independent experiments and four replicates each. The reporter gene activity of the *dctA* promotor (*dctAp*) fused to *lacZ* was also measured in a β-galactosidase assay. A volume of 250 μl per well was used for photometric measurements. In addition, cells were permeabilized by adding 200 μl of culture to 800 μl β-galactosidase buffer (KPi buffer (100 mM potassium phosphate, pH 7.0), potassium chloride (10 mM), magnesium chloride (1 mM), cetyltrimethylammonium bromide (0.005% w/v), sodium deoxycholate (0.0025% w/v)) with dithiothreitol (DTT, 8 mM). For the β-galactosidase assay 150 mL of permeabilized cells were transferred to 96 well plates and incubated at 30°C. To start the reaction 30 μl of ortho-nitrophenyl-β-D-galactoside (ONPG, 4 mg/mL) were added. The reaction was stopped after 20 min by addition of 70 μl Na_2_CO_3_ (1 M). Each strain was measured in two to four biological repeats and four independent samples each.

## Acknowledgements

We are grateful to Deutsche Forschungsgemeinschaft for funding (DFG UN 49/19-1 and UN 49/21-1). We thank Dr. Alexandra Kleefeld for the construction of parent reporter strains IMW385 and IMW386. We are also grateful to the National BioResource Project (NIG, Japan) for providing *E. coli* strains and to Dr. D. Ladant (Paris) for supplying the strains and plasmids for the BACTH system.

